# Information retrieval in an infodemic: the case of COVID-19 publications

**DOI:** 10.1101/2021.01.29.428847

**Authors:** Douglas Teodoro, Sohrab Ferdowsi, Nikolay Borissov, Elham Kashani, David Vicente Alvarez, Jenny Copara, Racha Gouareb, Nona Naderi, Poorya Amini

## Abstract

The COVID-19 pandemic has led to an exponential surge and an enormous amount of published literature, both accurate and inaccurate, a term usually coined as an infodemic. In the context of searching for COVID-19 related scientific literature, we present an information retrieval methodology for effectively finding relevant publications for different information needs. Our multi-stage information retrieval architecture combines probabilistic weighting models and re-ranking algorithms based on neural masked language models. The methodology was evaluated in the context of the TREC-COVID challenge, achieving competitive results with the top ranking teams participating in the competition. Particularly, the ranking combination of bag-of-words and language models significantly outperformed a BM25-based baseline model (16 percentage points for the NDCG@20 metric), correctly retrieving more than 16 out of the top 20 documents retrieved. The proposed pipeline could thus support the effective search and discovery of relevant information in the case of an infodemic.

## Introduction

In parallel to its public health crisis with vast social and economic impacts, the COVID-19 pandemic has resulted in an explosive surge of activities within scientific communities and across many disciplines^1^. The number of publications related to the pandemic has had an exponential growth since early 2020 when the pandemic was officially announced. In addition to the volume and velocity of the generated data, the heterogeneity of the data as a result of the typical variety of concept naming found in the biomedical field (e.g., COVID-19, SARS-COV2, Coronavirus disease 19, etc.), spelling mistakes (e.g., chloroquin, hydoxychloroquine), and the different source types (scientific reports, grey literature, preprint archives, etc.), among others, make searching and finding relevant literature within these corpora an important challenge.

To help clinical researchers, epidemiologists, medical practitioners and healthcare policy makers, among others, find the most relevant information for their needs, effective information retrieval models for these large and fast changing corpora became then a prominent necessity^2^. The information retrieval community, in turn, has responded actively and quickly to this extraordinary situation and has been aiming at addressing these challenges. To foster research for the scientific communities involved with the pandemic, the COVID-19 Open Research Dataset (CORD-19)^3^ collection was built to maintain all the related publications to the family of corona-viruses. This dataset helped research in various directions and several tasks are built around it, including natural language processing (NLP) related tasks, like question answering and language model pre-training, and information retrieval challenges in Kaggle^1^, as well as the TREC-COVID^2^.

The TREC-COVID^4, 5^ challenge ran in 5 rounds, each asking for an incremental set of information needs to be retrieved from publications of the CORD-19 collection. In a TREC-COVID round, participants were asked to rank documents of the CORD-19 corpus in decreasing order of likelihood of containing answers to a set of query topics. At the end of the round, experts provided relevance judgments for the top ranking documents submitted by different participants using a pooling strategy^4, 6^. Although limited to the first several top submissions of the participating teams, these relevance judgments are effective to evaluate the different models and are valuable training examples to train retrieval models for the subsequent rounds of the challenge.

More than 50 teams participated in the TREC-COVID challenge worldwide. Several of them have published their methodologies to tackle this complex task^7–12^. These works describe different information retrieval and NLP techniques as well as how to further adapt them to the infodemic case of the COVID-19. Having participated in the TREC-COVID challenge, we detail our retrieval methodology, which brought us competitive results with the top ranking teams. Particularly, we use a multi-stage retrieval pipeline, combining classic statistical weighting models with novel learning to rank approaches made by ensemble of deep masked language models. We present our results and analyse how the different components of the pipeline contribute to providing the best answers to the query topics.

## Related work

### Two-stage information retrieval

Currently, two main methodologies are used to rank documents in information retrieval systems: i) the classic query-document probabilistic approaches, such as BM25^13^ and probabilistic language models^14^, and ii) the learning-to-rank approaches, which usually post-process results provided by classic systems to improve the original ranked list^15, 16^. When there are sufficient training data, i.e., queries with relevance judgments for the case of information retrieval, learning-to-rank models often outperform classic one-stage retrieval systems^15, 17^. Nevertheless, empiric results have also shown that the re-ranking step may degrade the performance of the original rank^18^. Progress on learning-to-rank algorithms have been fostered thanks to the public release of annotated benchmark datasets, such as the LETOR^19^ and the Microsoft Machine Reading Comprehension (MS MARCO)^20^.

Learning-to-rank approaches can be categorised into three main classes of algorithms - pointwise, pairwise and listwise - based on whether they consider one document, a pair of documents or the whole ranking list in the learning loss function, respectively^15–17,21^. Variations of these learning-to-rank algorithms are available based on neural networks^16, 21^ and other learning algorithms, such as boosting trees^22^. More recently, pointwise methods leveraging the power of neural-based masked language models have attracted great attention^23, 24^. These learning-to-rank models use the query and document learning representations provided by the masked language model to classify whether a document in the ranked list is relevant to query. While these two-stage retrieval methods based on neural re-rankers provide interesting features, such as learned word proximity, in practice, the first stage based on classic probabilistic retrieval algorithms is indispensable, as the algorithmic complexity of the re-ranking methods makes them often prohibitive to classify the whole collection^17^.

Recent advances in text analytics, including question answering, text classification and information retrieval, have indeed mostly been driven by neural-based masked language models. A seminal effort in this direction is the Bidirectional Encoder Representations from Transformers (BERT) model^23^, which shows significant success in a wide range of NLP tasks. BERT uses a bi-directional learning approach based on the transformer architecture^25^ and is trained to predict masked words in a context. Since BERT was introduced, several works tried to augment its performance. A successful work in this direction is RoBERTa^26^, using larger and more diverse corpora for training as well as a different tokenizer. While RoBERTa needs larger computing power, it often improves the performance of BERT across different downstream tasks. Another similar effort is the XLNet model^27^, which uses a permutation-based masking, showing also consistent improvement over BERT.

### TREC-COVID retrieval efforts

Recently, the specific case of retrieval of COVID-related scientific publications has been addressed in several efforts^7–12^. These works follow mostly the above two-stage retrieval process. Among the first efforts is the SLEDGE system^7^, where the authors detail their solutions for the first round of TREC-COVID challenge using a BM25-based ranking method followed by a neural re-ranker. An important difficulty for the first round of the challenge is the absence of labelled data. To overcome this limitation, the authors lightly tune the hyper-parameters of the first stage ranking model using minimal human judgements on a subset of the topics. As for the second stage, they use the SciBERT^28^ model, which is pre-trained on biomedical texts, and fine-tuned on the general MS MARCO set^20^ with a simple cross-entropy loss. CO-Search^9^ uses a slightly different approach, wherein they incorporate semantic information, as captured by Siamese-BERT^29^, also within the initial retrieval stage. Moreover, they use the citation information of publications in their ranking pipeline. In the work of COVIDex^8^, the authors provide a full-stack search engine implementing a multi-stage ranking pipeline, where their first stage is based on the Anserini information retrieval toolkit^30^, complemented by different neural re-ranking strategies. They try to address the issue of length variability among documents with and atomic document representation using, for example, paragraph-level indexing.

## Methods

In this section, we describe the corpus and query set, and our methodology for searching COVID-19 related literature in the context of the TREC-COVID challenge. We start by introducing the CORD-19 dataset, which is the corpus used in the competition. We then describe the challenge organisation and assessment queries. Then, we detail our searching methodology, based on multi-stage retrieval approach. Finally, we present the evaluation criteria used to score the participant’s submissions. For further details on the TREC-COVID challenge, please see^4, 5^.

### The CORD-19 dataset

A prominent effort to gather publications, preprints and reports related to the corona-viruses (COVID-19, SARS, MERS) is the CORD-19 collection of the Allen Institute for Artificial Intelligence (in collaboration with other partners)^3^. Figure 1 describes the size and content origin of the corpus for the different TREC-COVID rounds. As we can see, this is a large and dynamically growing semi-structured dataset from various sources like PubMed, PubMed Central, WHO and preprint servers like bioRxiv, medRxiv, and arXiv. The dataset contains document metadata, including *title, abstract, authors*, among others, but also the full text or link to full text files when available. A diverse set of related disciplines, e.g., from virology and immunology to genetics, are represented in the collection. Throughout the challenge, the dataset was updated on a daily basis and snapshot versions representing its status at a certain time were provided to the participants for each round. In the last round of the TREC-COVID challenge, the corpus contained around 200’000 documents, coming mostly from Medline, PMC and WHO sources.

**Figure 1.**
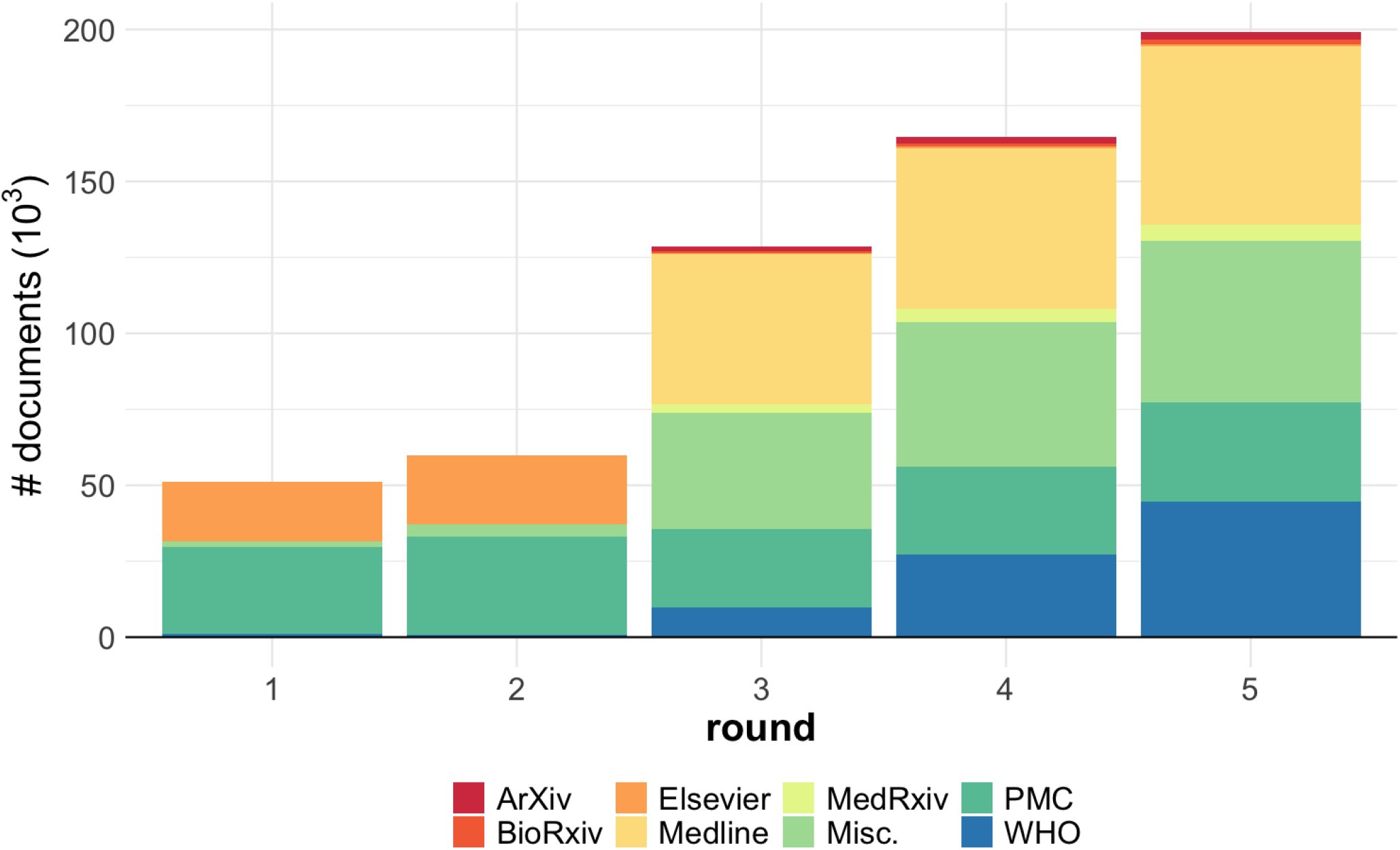
Evolution of the CORD-19 corpus across the TREC-COVID rounds stratified by source.

### The TREC-COVID challenge

To assess the different information retrieval models, the TREC-COVID challenge provided a query set capturing important information search needs of researchers during the pandemic^4, 5^. These needs are stated in different query topics, consisting of three free text fields - *query, question* and *narrative* - with an increasing level of context, as shown in the example of Table 1. The challenge started with 30 topics in round 1 and added five new topics at each new round, reaching thus 50 topics in round 5.

In each round, the participants provided ranked lists of candidate publications of the CORD-19 collection, which best answered the query topics. Each list was generated by a different information retrieval model, so called *run*, with up to 5 runs in the first 4 rounds and 7 runs in the last round per team. At the end of the round, domain experts examined the top k candidate publications (where k is defined by the organisers) from the priority runs of the teams and judged them as “highly relevant”, “somehow relevant”, or “irrelevant”. Then, based on the consolidated relevance judgments, the participants were evaluated using standard information retrieval metrics (NDCG, precision, etc.). Judged documents for a specific topic from previous rounds were excluded from the relevance judgement list.

**Table 1.** Example of a TREC-COVID topic (# 1 out of 50) with the fields *query, question* and *narrative*.)

|  |  |
| --- | --- |
| **topic** | 1 |
| **query** | coronavirus origin |
| **question** | what is the origin of COVID-19 |
| **narrative** | seeking range of information about the SARS-CoV-2 virus's origin, including its evolution, animal source, and first transmission into humans |

### Proposed retrieval methodology

Figure 2 shows the different components of our information retrieval pipeline for the COVID-related literature. These components can be divided into three main categories: *i)* first-stage retrieval using classic probabilistic methods; *ii)* second-stage (neural) re-ranking models; and *iii)* rank fusion algorithms. Given a corpus containing metadata information, such as title and abstract, and full text, when available, documents are stored using directed and inverted indexes. Then, transformer-based and classic learning-to-rank models trained using relevance judgements are used to classify and re-rank pairs of query-document answers. The ranked list obtained from the different models are further combined using the reciprocal rank fusion (RRF) algorithm.

**Figure 2.**
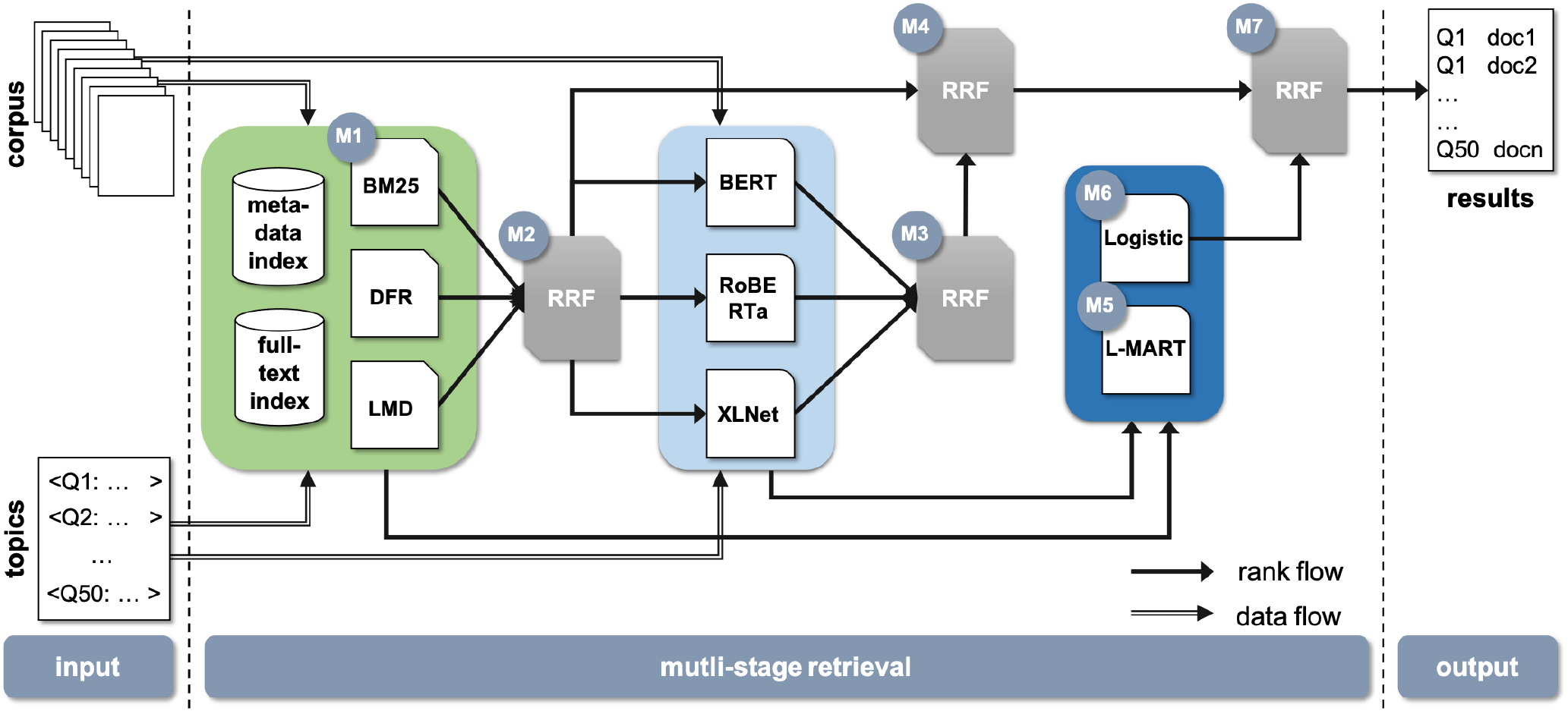
Multi-stage retrieval pipeline. Light green: first-stage retrieval. Light and dark blue: second-stage retrieval. M1-M7 denote the different models to create the respective runs 1-7 in round 5. RRF: Reciprocal Rank Fusion. Logistic: logistic regression model. L-MART: LambdaMART model.

#### First-stage retrieval

For the first-stage retrieval, we assessed three variations of the classic query-document probabilistic weighting models: Okapi Best Match 25 (BM25)^31^, Divergence from Randomness (DFR)^32^ and Language Model Dirichlet (LMD)^33^.

### Okapi Best Match 25

Our first classical model, Okapi BM25^31^, is based on the popular term frequency-inverse document frequency (tf-idf) framework. In the tf-idf framework, term weights are calculated using the product of within term-frequency *t f* and the inverse document frequency *id f* statistics. Denote *f* (*t, d*) as the number of times a term *t* appears in a document *d* within a collection *D*, BM25 calculates the term-weight *w* as:

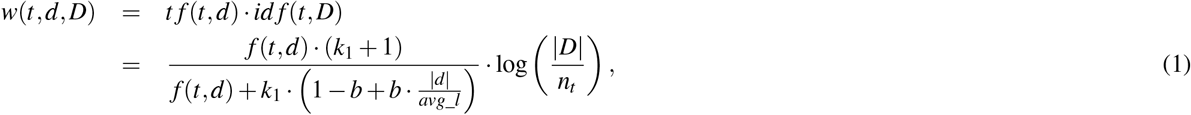

where |*d*| is the length of the document, |*D*| is the size of the collection, *avg*_*l* is the average length of the documents in the collection, *n*_*t*_ is the number of documents containing the term *t*, and *k*_1_ and *b* are parameters of the model associated with the term frequency and the document size normalisation, respectively.

### Divergence from Randomness

The second model, DFR, extends the basic tf-idf concept by considering that the more the divergence of the term-frequency *t f* from its collection frequency *c f* (*c f*≈ *d f*), the more the information carried by the term in the document^32^. Thus, for a given model of randomness *M*, in the DFR framework the term-weight is inversely proportional to the probability of term-frequency within the document obtained by *M* for the collection *D*:

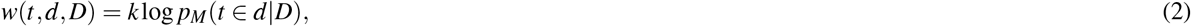

where *p*_*M*_ is a probabilistic model, such as binomial or geometric distributions, and *k* a parameter of the probabilistic model.

#### Language Model Dirichlet

The third model, LMD, uses a language model, which assigns probabilities to word sequences, with a Dirichlet-prior smoothing, to measure the similarity between a query and a document^33^. In a retrieval context, a language model specifies the probability that a document is generated a query, and smoothing is used to avoid zero probabilities to unseen words and improves the overall word probability accuracy. In the LMD, term-weigh is calculated using the following equation:

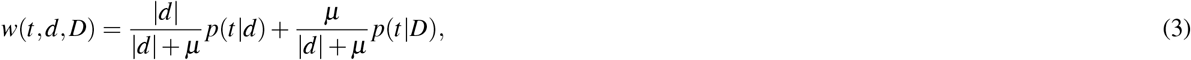

where *p*(*t* |*d*) denotes the probability of a term in a document, *p*(*t* |*D*) is the probability of a term in the collection, and *µ* is the Dirichlet parameter to control the amount of smoothing.

In our pipeline, the BM25, DFR and LMD implementations are based on the Elasticsearch framework^3^. The model parameters were trained using the relevance judgements of the round 4 in a 5-fold cross-validation setup.

#### Second-stage re-ranking

The models used in the first-stage ranking are based on the bag-of-words statistics, where essentially we look at the histogram of query terms, and their document and collection statistics but neglect the sequential nature of text and word relations. After the first-stage retrieval, we use neural masked language models trained on the relevance judgements from previous rounds to take into account the sequential nature of text and improve the initial rankings. As shown in Figure 2, we assessed three masked language models based on the transformer architecture: BERT, RoBERTa and XLNet.

### BERT

Figure 3 shows the general idea of how we use the BERT language model to match documents to a query topic. Given a topic and a document associated to it as input and a relevance judgement as the label for the query-document association (relevant or not), the model is trained or fine-tuned in the BERTology parlance, as it had been previously pre-trained on a large corpus, to predict whether the document is relevant or not to the query. In the input layer of the pre-trained model, the topic and candidate publication are tokenized and separated by the language model `[SEP]` token (stands for sentence separation). Moreover, to enforce the sequential structure of text, positional embedding as well as sentence embedding are added to the main embeddings for each token. These embeddings are then fed to the transformer layers of BERT, which are updated during the fine-tuning step. Finally, the output of the special `[CLS]` token (stands for classification) is used to determine the relevance of the candidate publication to the queried information topic.

**Figure 3.**
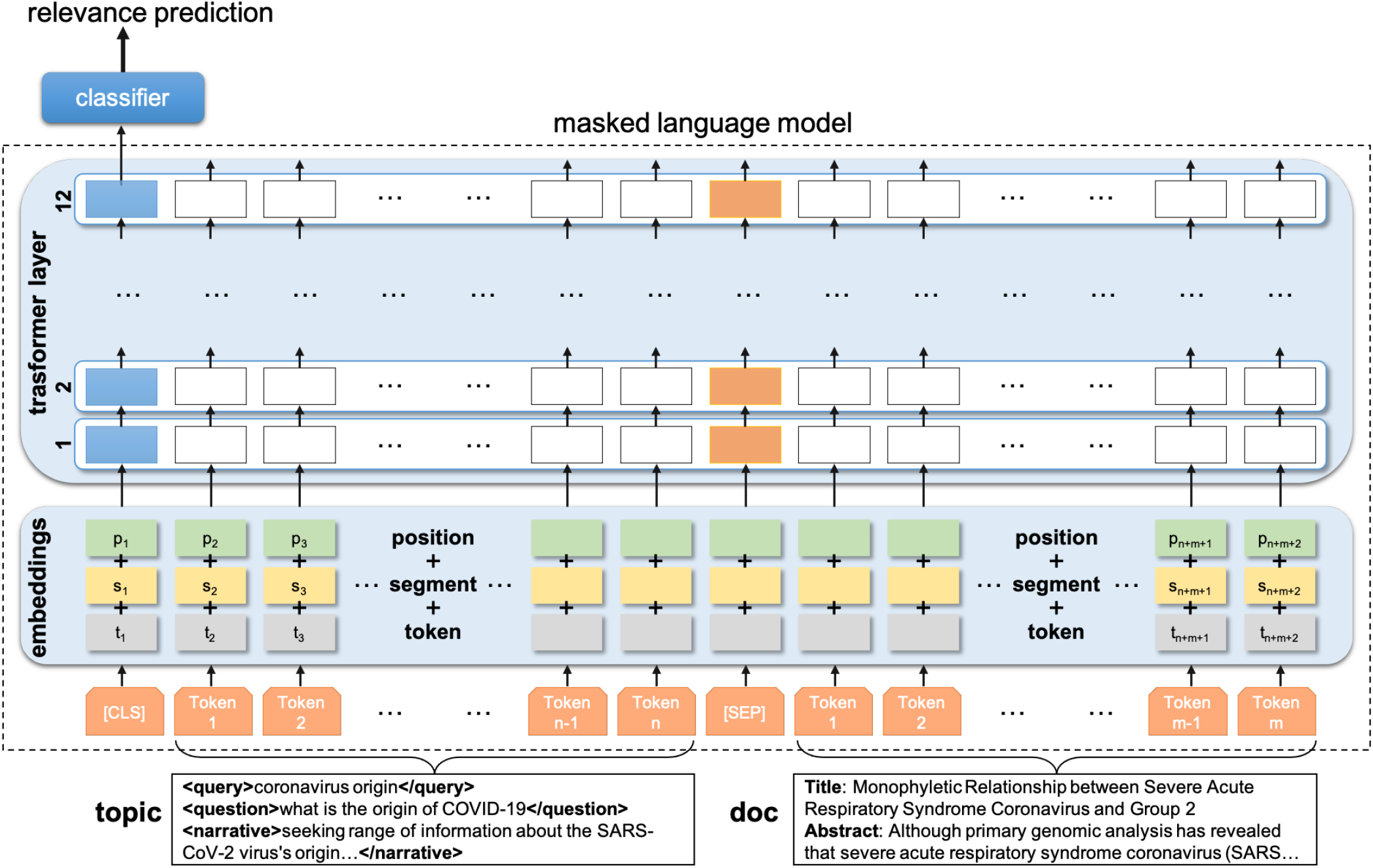
Neural masked language model for document relevance classification. As inputs to the pre-trained masked language model, the topics and candidate publications are separated by the `[SEP]` tag. Inputs are tokenized using sub-words tokenization methods (gray boxes). Segment embeddings (yellow boxes) represent the difference between a topic and a document input. Position embeddings (green boxes) enforce the sequential structure of text. The transformer and classification layers are updated in the training phase using the relevance judgements. The output of the special `[CLS]` token is finally used to determine the relevance of the candidate publication to the queried information topic.

Using the query topics from a preceding round (round 4 for the results presented here) and their respective list of relevance judgments, we fine-tuned the BERT model to re-score the initial association of the query-document pair between 0 (not relevant) and 1 (very relevant). For this, we use the score associated to the `[CLS]` token position. We limit the input size of the query and document to 512 tokens (or sub-words). Then, at the second-stage re-ranking step we classify the top *k* publications retrieved by the first stage models using the fine-tuned BERT model (we set *k* = 5000 in our experiments).

### RoBERTa and XLNet

Identical training strategies were used for the RoBERTa and XLNet language models. The main difference for the RoBERTa model is that it was originally pre-trained on a corpus with an order of magnitude bigger compared to BERT (160GB *vs*. 16GB). Moreover, it uses dynamic masking during the training process, that is, at each training epoch, the model sees different versions of the same sentence with masks on different positions, compared to a static mask algorithm for BERT. Lastly, RoBERTa uses a byte-level Byte-Pair-Encoding tokenizer compared to BERT’s WordPiece. As BERT and its variants (e.g., RoBERTa) neglect the dependency between the masked positions and suffer from a pretrain-finetune discrepancy, XLNet adopts a permutation language model instead of masked language model to solve the discrepancy problem. For downstream tasks, the fine-tuning procedure of XLNet is similar to that of BERT and RoBERTa.

We use the BERT, RoBERTa and XLNet model implementations available from the HuggingFace framework^4^. The models were trained using the Adam optimiser with an initial learning rate of 1.5*e −* 5, weight decay of 0.01, and early stopping with a patience of 5 epochs.

#### Combining model results

We use the RRF algorithm^34^ to combine the results of different retrieval runs. RRF is a simple, yet effective technique to re-score the retrieval list based on the scores of multiple retrieval lists. Given a set *D* of documents to be sorted and a set *R* = *{r*_1_…*r*_*n*_*}* of ranking files, each with a permutation on 1…|*D*|, RRF computes the aggregated score using the following equation:

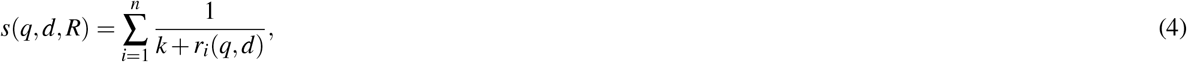

where *r*(*q, d*) is the rank of document *d* for the query *q* in the ranking file *r*_*i*_ and *k* is a threshold parameter, which was tuned to *k* = 60 using previous round’s data.

#### Second-step learning-to-rank

To exploit the features (relevance score) created by the different bag-of-words and masked language models, we added a second-step learning-to-rank pass to our pipeline. Using the similarity scores *s* computed by the BM25, DFR, LMD, BERT, RoBERTa and XLNet as input features and the relevance judgements of previous rounds as labels, we trained two learning-to-rank models: LambdaMART and a logistic regression classifier. While the language models exploit the sequential nature of text, they completely neglect the ranking provided by the bag-of-words models. Thus, we investigate the use of the LambdaMART^16^ algorithm, which uses a pairwise loss that compares pairs of documents and tells which document is better in the given pair. Moreover, we trained a simple pointwise logistic regression that takes into account the ranks extracted by fist- and second-stage retrieval models. We used the pyltr^5^ and scikit-learn^6^ implementations for the LambdaMART and logistic regression, respectively. For the LambdaMART model we trained the learning rate and the number of estimators, and for the logistic regressions we trained the solver and regularization strength parameters.

#### First-stage retrieval: pre-processing, querying strategies and model fine-tuning

In the first-stage retrieval step, we apply a classical NLP pre-processing pipeline to the publications (indexing phase) and topics (search phase): lower-casing, removal of non alphanumerical characters (apart from “-”), Porter stemming. Additionally, a minimal set of COVID-related synonyms, such as “covid-19” and “sars-cov-2”, were created and used for query expansion.

The queries are then submitted to the index in a combinatorial way using the different topic fields and document sections. This means that, for each of the *query, question* and *narrative* fields of a topic, we submit a query against the index for each of the *title* and *abstract* sections of the publications (*abstract* + *full text* in case of the full text index). Additionally, the whole topic (query + question + narrative) was queried against the whole document. This querying strategy leads to 7 queries for each topic and the final score is computed by summing up the individual scores. Moreover, as the first publicly announce of a coronavirus-related pneumonia was made in January 2020, we filter out all publications before December 2019.

We define the best query strategy and fine-tune the basic parameters of the bag-of-words models using the relevance judgements of the previous round in a 5-fold cross-validation approach. As an example, to tune the *b* and *k* parameters of the BM25 model at round 5, we take the topics and relevance judgement of round 4 and submit to the index of round 5, optimizing the P@10 metric. For round 1, we used default parameter values.

### Evaluation criteria

We use the official metrics of the TREC-COVID challenge to report our results: precision at K documents (P@K), normalized discounted cumulative gain at K documents (NDCG@K), mean average precision (MAP) and binary preference (Bpref)^4^. For all these metrics, the closest to 1, the best is the retrieval model. They are obtained using the the trec_eval information retrieval evaluation toolkit^7^.

## Results

Seven models of our pipeline were used to create the 7 runs submitted for the official evaluation of the TREC-COVID challenge (labels M1 to M7 in Figure 2). Our first model - *bm25* - based on the BM25 weighting model against the metadata index provides the baseline run. Our second model - *bow + rrf* - is a fusion of the BM25, DFR and LMD weighting models computed against the metadata and full text indices and combined using the RRF algorithm. Model 3 - *mlm + rrf* - uses the RRF combination of BERT, RoBERTa and XLNet models applied to the top 5000 documents retrieved by model 2. Model 4 - *bow + mlm + rrf* - combines the results of model 2 and 3 using the RRF algorithm. Then, model 5 - *bow + mlm + lm* - re-ranks the results of runs 2 and 3 using the LambdaMART algorithm trained using the similarity scores of the individual models 2 and 3. Similarly, model 6 - *bow + mlm + lr* - is based on a logistic regression classifier that uses as classification features for the query-document pairs the similarity scores of runs 2 and 3. Finally, model 7 - *bow + mlm + lr + rrf* - combines runs 2, 3 and 6 using the RRF algorithm. For all RRF combinations, the parameter *k* was set to 60. All models and parameters were trained using round 4 relevance judgments. Table 2 summarizes the submitted runs.

**Table 2.** Summary of the submitted runs. Refer to Figure 2 for a pictorial description.

| run | name | description |
| --- | --- | --- |
| 1 | bm25 | Run based on the baseline BM25 model using the metadata index. |
| 2 | bow + rrf | A RRF combination of BM25, DFR and LMD models computed against the metadata and full text indices. |
| 3 | mlm + rrf | A RRF combination of BERT, RoBERTa and XLNet models applied to run 2. |
| 4 | bow + mlm + rrf | A RRF combination of runs 2 and 3. |
| 5 | bow + mlm + lm | A LambdaMART-based model using features from the individual models used to create runs 2 and 3. |
| 6 | bow + mlm + lr | A logistic regression model using features from the individual models used to create runs 2 and 3. |
| 7 | bow + mlm + lr + rrf | A RRF combination of runs 2, 3 and 6. |

### Official evaluation results

Table 3 shows the official results of the TREC-COVID challenge for the 7 submitted runs. As we can see, the best results are provided by model 7 (bow + mlm + lr + rrf), apart from the metric Bpref, which is the highest for model 5 (bow + mlm + lm). Comparing the NDCG@20 metric, model 7 improved 16.4 percentage point against to the baseline model (26.0% of relative improvement). Overall, almost 17 out the top 20 documents retrieved by model 7 were pertinent to the query. Model 3 was able to retrieve 6.6% more relevant documents compared to the baseline model (6963 *vs*. 6533 out of a total of 10910 documents judged relevant for the 50 queries). On the other hand, it showed a relative improvement in precision at the top 20 documents of 22.4%. Therefore, it not only improved the recall but also brought relevant documents higher in the ranking list. These results show that the use of the masked language models had a significant positive impact in the ranking.

**Table 3.**
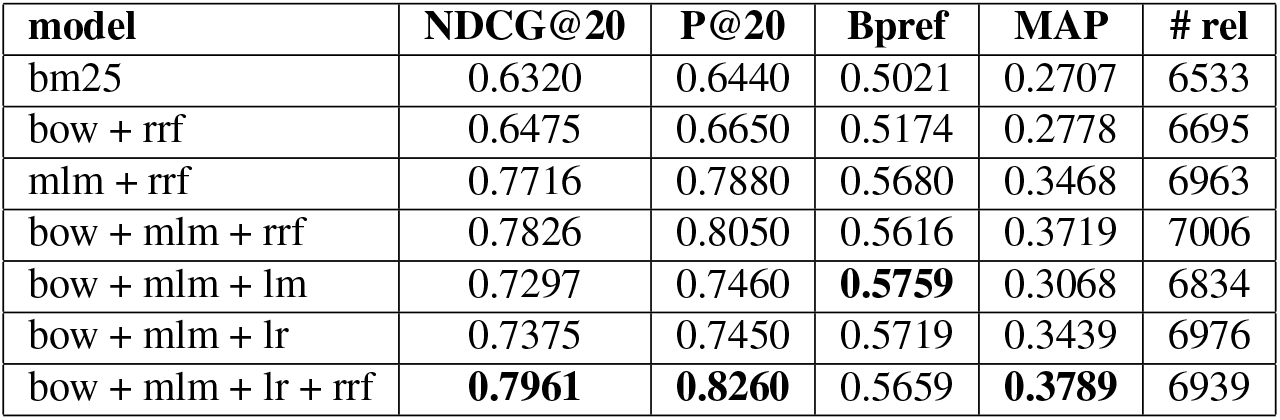
Performance of our models in round 5 of the TREC-COVID challenge. # rel is the total number of relevant documents retrieved by the model for the 50 queries. A total of 10910 documents were judged relevant by the organisers.

| model | NDCG@20 | P@20 | Bpref | MAP | # rel |
| --- | --- | --- | --- | --- | --- |
| bm25 | 0.6320 | 0.6440 | 0.5021 | 0.2707 | 6533 |
| bow + rrf | 0.6475 | 0.6650 | 0.5174 | 0.2778 | 6695 |
| mlm + rrf | 0.7716 | 0.7880 | 0.5680 | 0.3468 | 6963 |
| bow + mlm + rrf | 0.7826 | 0.8050 | 0.5616 | 0.3719 | 7006 |
| bow + mlm + lm | 0.7297 | 0.7460 | **0.5759** | 0.3068 | 6834 |
| bow + mlm + lr | 0.7375 | 0.7450 | 0.5719 | 0.3439 | 6976 |
| bow + mlm + lr + rrf | **0.7961** | **0.8260** | 0.5659 | **0.3789** | 6939 |

Table 4 shows the official best results for the different metrics for the top 10 teams participating in round 5 of TREC-COVID (NDCG@20 metric taken as reference). Comparing the NDCG@20 metric, the best model submitted by our team (risklick) were ranked 4 out of the 28 teams participating in round 5, 5.4 percentage point below the top performing team (Unique-ptr). For a reference, the best performing model in the challenge retrieves on average 17.5 relevant documents per query in the top 20 retrieved documents compared to 16.5 for our model. If we consider a reference baseline made by the median of the participating teams’ best value, our pipeline outperforms the baseline by 11.7%, 14.6%, 16.7% and 25.0% for the MAP, P@20, NDCG@20, and Bpref metrics, respectively. All data and results of the TREC-COVID challenge can be found at: https://ir.nist.gov/covidSubmit/data.html.

**Table 4.**
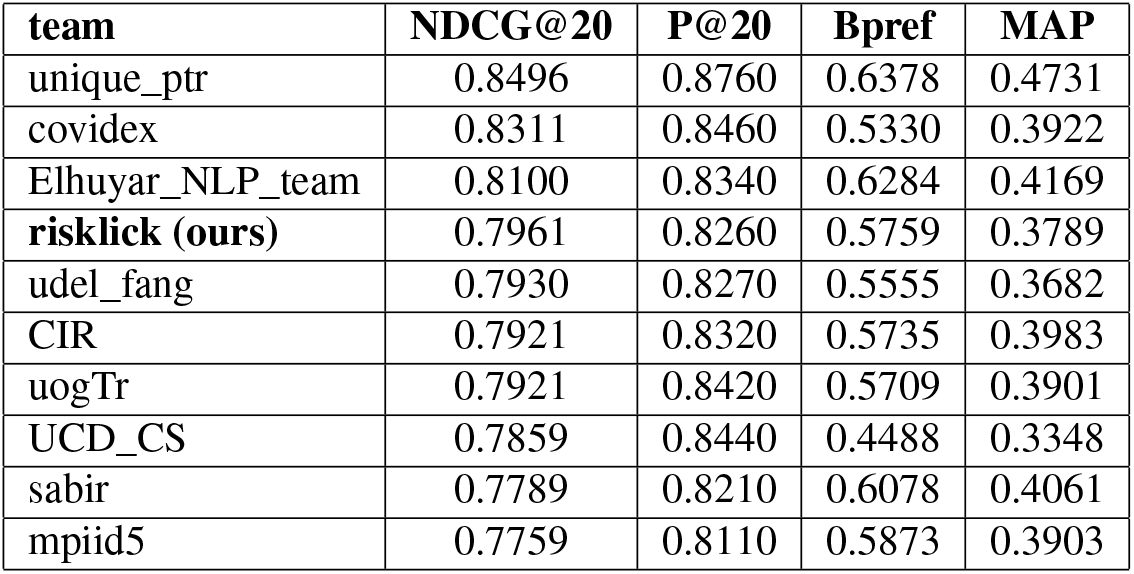
Official leaderboard for the final round of the TREC-COVID challenge. Best results for the top 10 teams using the NDCG@20 as reference metric. A total of 28 teams participated in TREC-COVID final round.

### Model performance analyses

Figure 4 shows the relative improvement of the different models in the pipeline in relation to the baseline (model 1 - bm25) according to the NDCG@20 metric. The most significant contribution to the final performance comes from the inclusion of the masked language models in the pipeline (model 3 - mlm + rrf), adding a relative performance gain of 22.4% to the results. The learning-to-rank models based on classic machine learning models - model 5 and model 6 - actually jeopardise the performance when compared to their previous model in the pipeline (model 4). However, when model 6 is combined with model 4, a 2.1 percentage point gain is achieved on top of model 4, leading to the best model (model 7 - bow + mlm + lr + rrf). Indeed, it is important to notice the consistent benefit of combining models using the RRF algorithm. Interestingly, the effect of LambdaMART seems to be significantly detrimental for *P*@20, *NDCG*@20 and *MAP*, but marginally beneficial for *Bpref*, for which it is the best model.

**Figure 4.**
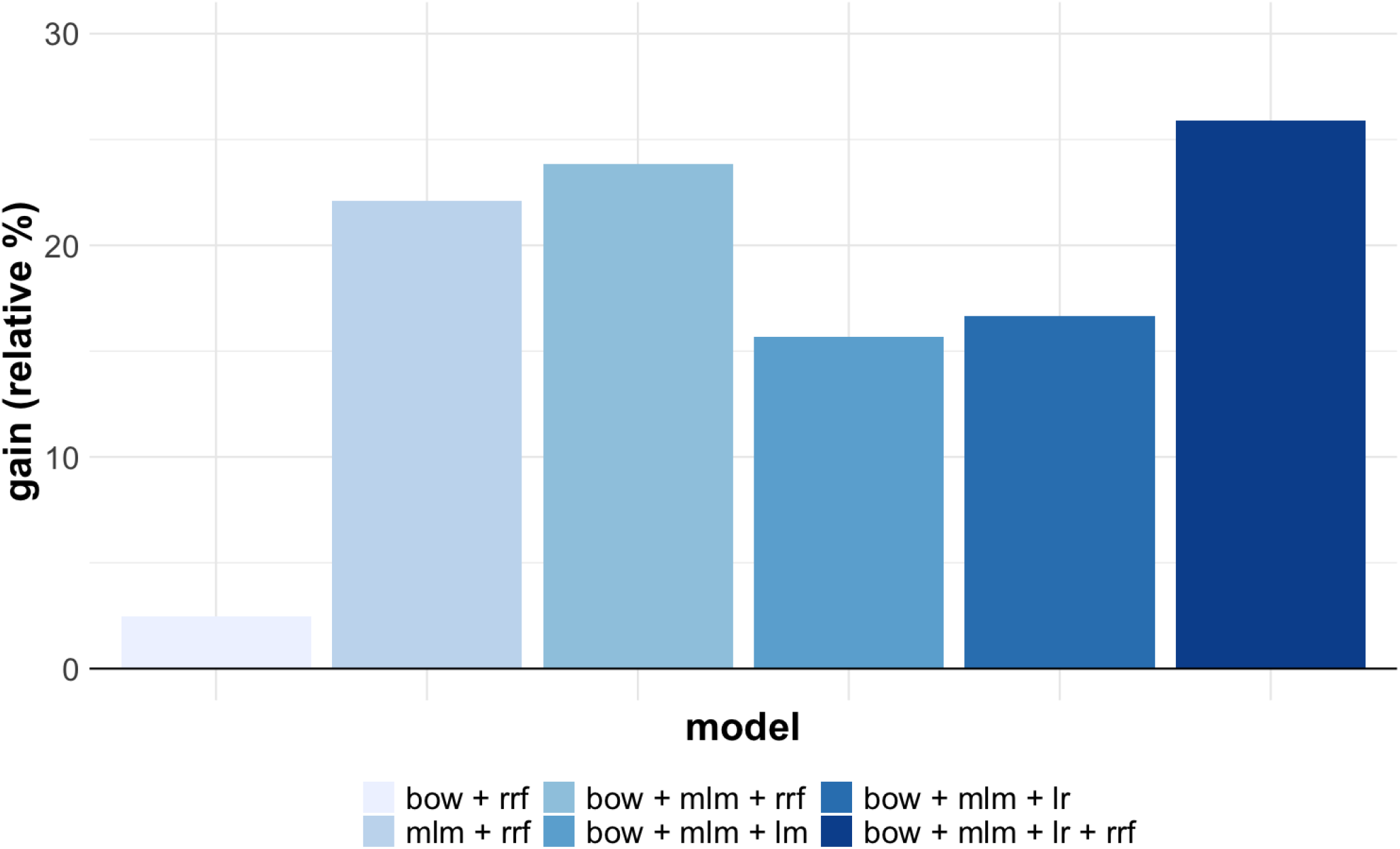
Relative contribution of each model for the *NDCG*@20 metric compared to the baseline model bm25.

The performance of the individual masked language models is shown in Table 5. Surprisingly, they are similar to the baseline model, with small performance reductions for BERT and RoBERTa models, and a small performance gain for the XLNet model. However, when combined they provide the significant performance improvement shown in Figure 4. Our assumption is that they retrieve different documents as relevant and their combination using RRF ends up aligning these documents in the top rank. Indeed, looking at the top 3 documents for query 1 retrieved by these models, for example, there is no overlap between the documents, being 8 relevant and 1 unjudged. This result clearly shows the beneficial effect of using ensemble of masked language models, as well as the success of RRF in fusing their retrievals.

**Table 5.** Performance of the individual masked language models and their combination using RRF.

| model | NDCG@20 | P@20 | Bpref | MAP | # rel |
| --- | --- | --- | --- | --- | --- |
| BERT | 0.6209 | 0.6430 | 0.5588 | 0.2897 | 6879 |
| RoBERTa | 0.6261 | 0.6440 | 0.5530 | 0.2946 | 6945 |
| XLNet | 0.6436 | 0.6570 | 0.5644 | 0.3064 | 6926 |
| mlm + rrf | 0.7716 | 0.7880 | 0.5680 | 0.3468 | 6963 |

### Topic performance analyses

The performance analyses for the individual topics shows that our best model has a median of 0.9000 for the P@20 metric (max=1.0000; min=0.3000), which demonstrates an effective overall performance. However, as shown in Figure 5, for some topics, notably 11, 12, 19, 33 and 50, less than 50% of documents in the top 20 retrieved are relevant. For topics 11, 12, and 19, which searches for *coronavirus hospital rationing, coronavirus quarantine* and *what alcohol sanitizer kills coronavirus* information, respectively, all of our models have a poor performance and indeed the combination of the different models in the pipeline manages to boost the results. On the other hand, for topics 33 and 50, which searches for *coronavirus vaccine candidates* and *mRNA vaccine coronavirus* information, respectively, it was the combination with the logistic regression model that lowered the performance (notice in Figure 5 that model 4 - bow + mlm + rrf has a significant better compared to model 7 for those queries).

**Figure 5.**
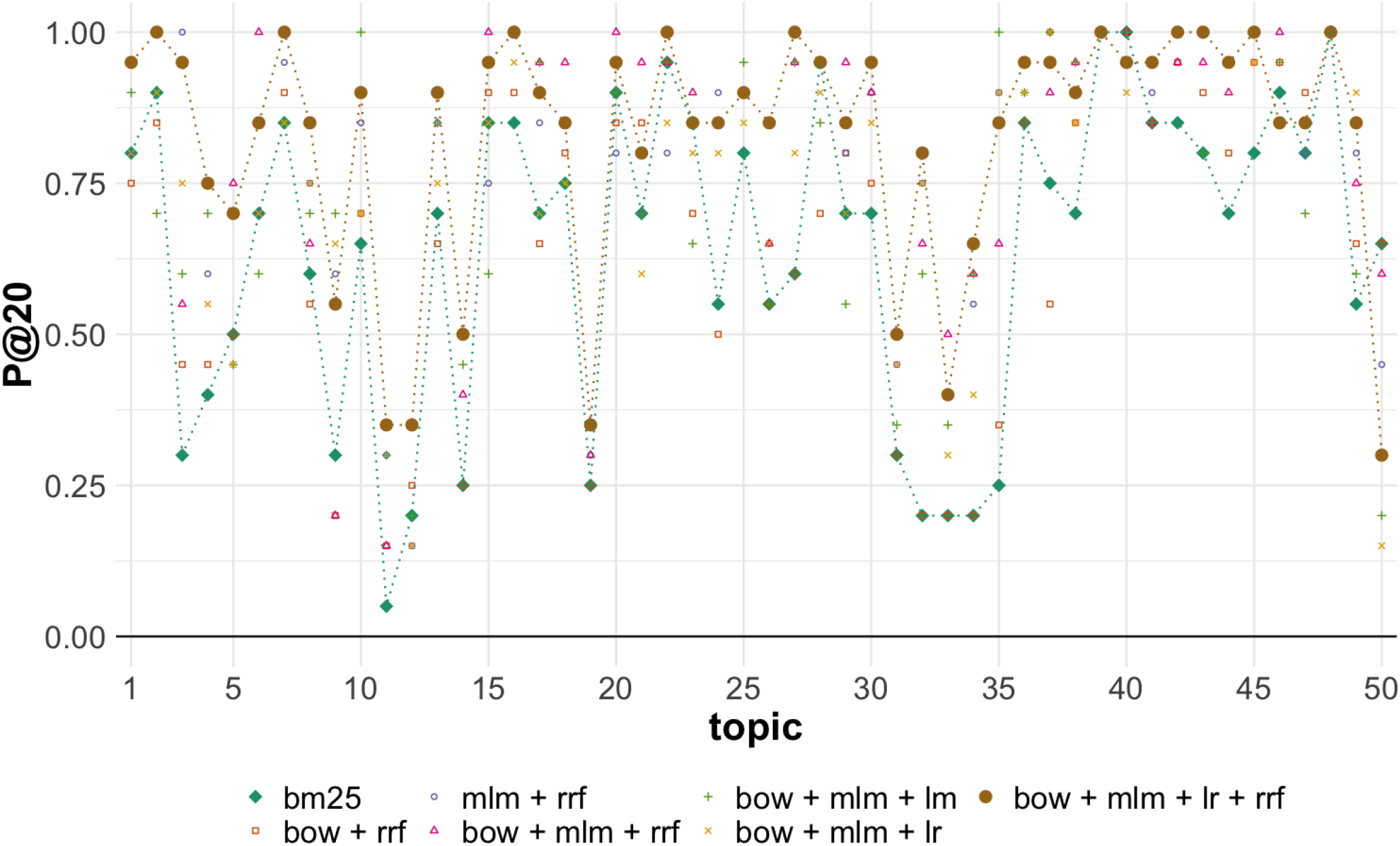
Per topic number of relevant publications retrieved up to rank 50 of round 5 of TREC-COVID per each run. The baseline run1 and the best-performing run7 which benefits from neural language models are highlighted with dashed lines. Note that for most topics, the transformer-based runs have significantly improved performance.

The difference in performance per topic between our best model and the median of the submitted runs in round 5 for all teams for the P@20 metric is shown in Figure 6. Indeed, topics 11, 12 and 19 seem hard for all the models participating in the TREC-COVID challenge to retrieve the correct documents. Even if our best model has a poor performance for them, it still outperforms most of the runs submitted to the official evaluation. In particular, topic 19 has only 9 relevant or somewhat relevant documents in the official relevance judgements, which makes that its max performance be slightly less than 50% for the P@20 metric. For our worst performing topics compared to the other participants - topics 33 and 50 - a better tuning between the ranking weights of the bag-of-words, masked language and logistic regression models could have boosted the results.

**Figure 6.**
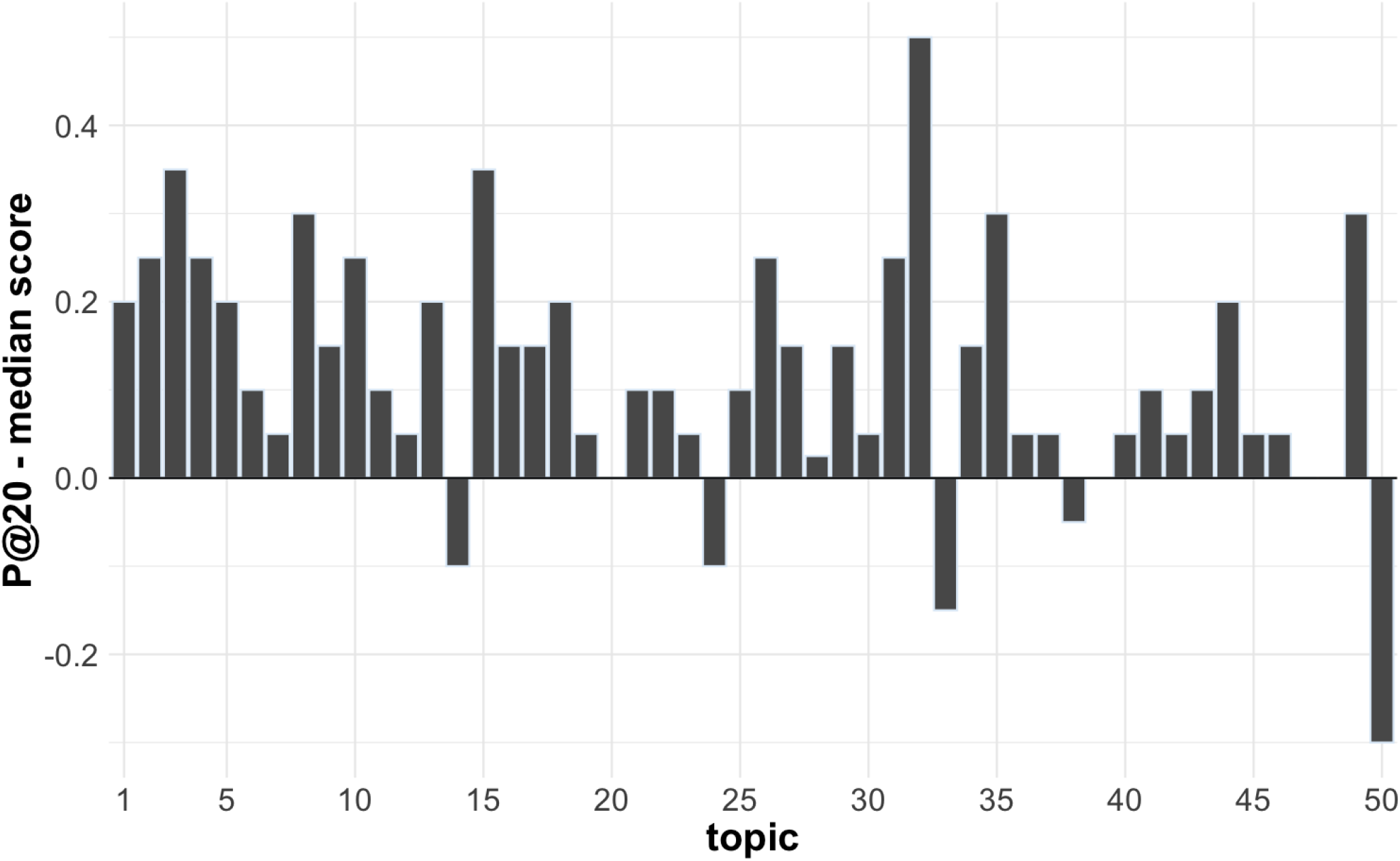
Per topic performance difference between our best model (model 7) and the median of all official submissions for the P@20 metric in round 5.

### Time-dependent relevance analyses

Given the dynamics of the COVID-19 pandemics, with a starting period relatively well defined, a particularly effective technique to remove noise from the results, also adopted by some other participating teams^7^, is filtering documents based on their publication dates. For our first-stage retrieval models, we filtered out publications before December 2019, when the outbreak was first detected. This led to small impact on recall but highly improved the precision of our models.

To better understand how the document relevance varied over time, we analysed the publication date of the official relevance judgements for the five rounds of TREC-COVID. As we can see in Figure 7, there is a clear decay pattern in the number of relevant articles over time for all the rounds, with a faster decay in the first rounds and a longer tail for the later ones. We notice that more recent publications, closer to round start when the snapshot of the collection was created and queries were submitted, tend to have a higher probability of being relevant to the information need. This is somehow expected as the documents found in previous query rounds were explored and are no longer relevant, only the most recent data is interesting, particularly in the gap between rounds. A second explanation is that in the case of a pandemic, new evidence arrives with an explosive rate, possibly refuting older knowledge.

**Figure 7.**
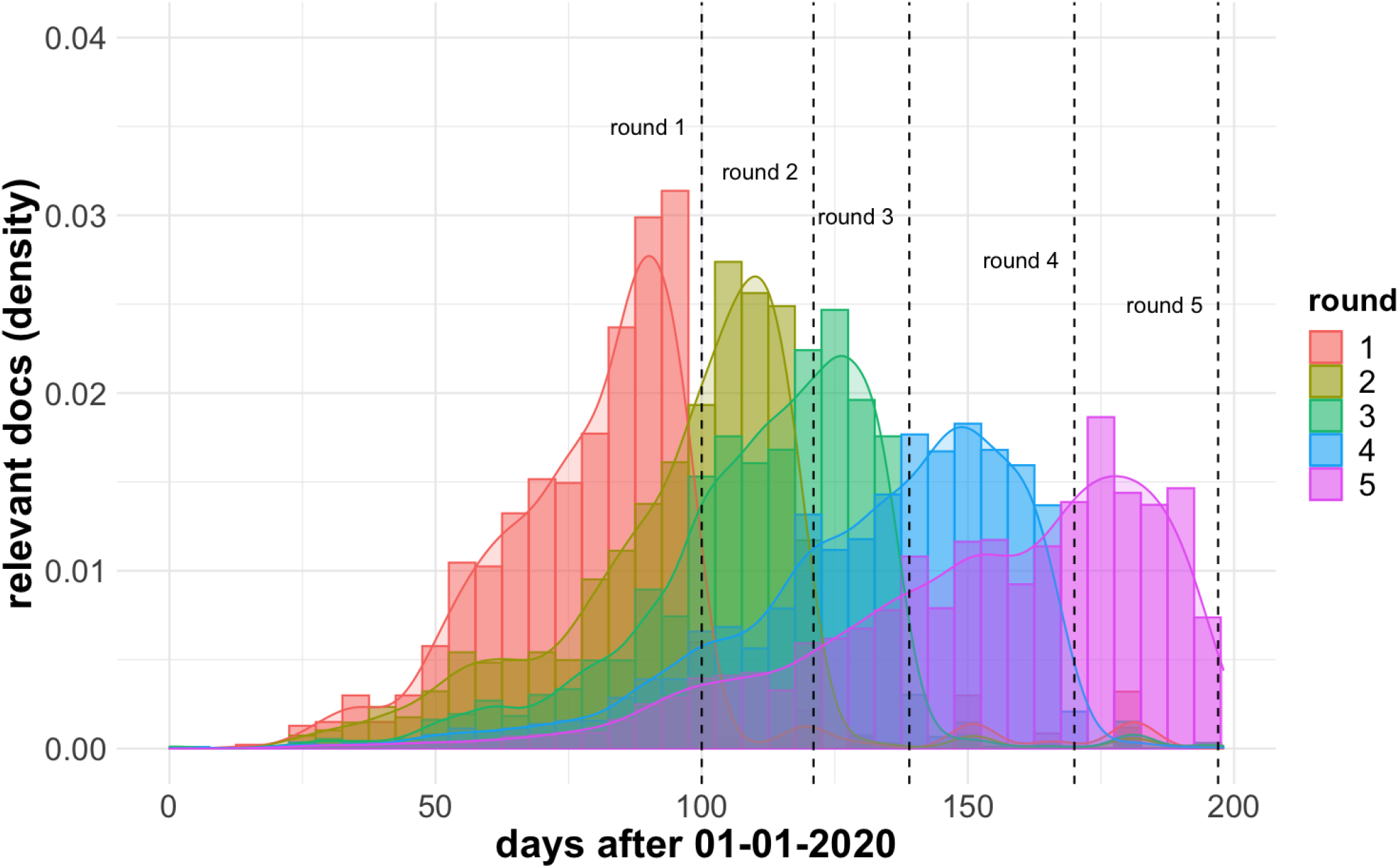
Distribution of the publication dates of the “highly relevant” articles for each of the TREC-COVID rounds.

## Discussion

The COVID-19 pandemic has led to huge amount of literature being published in most diverse sources, including scientific journals, grey repositories, white reports, among others. As the pandemic continues, the number of scientific publications grows in an unprecedented rate causing an infodemic within many different disciplines involved. Finding the most relevant information sources to answer different information needs within the huge volume of data created becomes of utmost necessity. To support effective search for COVID-19-related information, we explored the use of a multi-stage retrieval pipeline supported by bag-of-words models, masked language models and classic learning-to-rank methods. The proposed methodology was evaluated in the context of the TREC-COVID challenge and achieved competitive results, being ranked top 4 out of 126 runs from 28 teams. The use of the multi-stage retrieval approach significantly improved the results, leading to a gain in performance of 25.9% in terms of the NDCG@20 metric compared to a bag-of-words baseline. The ensemble of masked language models brought the highest performance gain to the pipeline. Indeed, ensemble of language models has proved to be a robust methodology to improve predictive performance^35–37^.

Looking at the boost in performance alone, one could be tempted to argue that masked language models could be the main component in a retrieval system. However, two issues may arise: algorithmic complexity and effectiveness. The former is related to the high complexity of masked language models (*𝒪*(*n*^2^ *· h*), where *n* is the sentence length and *h* is the number of attention heads), which makes it prohibitive to classify a whole collection, often containing millions of documents, for a given query. The latter is related to the effectiveness of the individual models themselves. As shown in Table 5, alone the performance of the individual language models is not significantly different from the baseline BM25 model. Thus, we believe it is the effective combination of models with different properties that can provide a successful search strategy in complex corpora, as the one originated from the COVID-19 infodemic.

With the rapid surge of published information, and the variety of topics and sources related to COVID-19, it became hard for professionals dealing with the pandemic to find the correct information for their needs. While the automation discussed in this work can support more effective search and discovery, some high level topics are still challenging. Indeed, some topics assessed in the TREC-COVID challenge showed to be particularly hard to the retrieval models. For example, for topic 11, which searched for documents providing information on “guidelines for triaging patients infected with coronavirus”, our best model prioritized documents providing information about indicators for diagnosis (“early recognition of coronavirus”, “RT-PCR testing of SARS-CoV-2 for hospitalized patients clinically diagnosed”, etc.). On the other hand, it missed documents including passages such as “telephone triage of patients with respiratory complaints”. Similarly, for topic 12, which searched information about the “best practices in hospitals and at home in maintaining quarantine”, our model prioritized documents providing information about “hospital preparedness” (“improving preparedness for”, “preparedness among hospitals”, etc.) and missed documents containing information about “home-based exercise note in Covid-19 quarantine situation”.

The TREC-COVID challenge dynamics, running throughout a sequence of rounds with new incremental search topics added on each round provides an interesting setting for evaluating retrieval models in an infodemic context. It simulates typical search and discovery workflows, in which evolving queries are posed against an evolving body of knowledge over time, and already discovered documents in previous searches are no longer relevant^38, 39^. An effective strategy in this case is to filter out results according to a cut-off date, reducing thus noise in the retrieval set. However, in retrospect, we notice that another useful technique, which is very natural to an infodemic case, could be to decay the score of publications by their distance to the present time or explore their recency or freshness^40, 41^, as highlighted in Figure 7, rather than a hard cut-off (i.e., December 2019 in our case). We leave exploring such strategy as future work.

To conclude, we believe our information retrieval pipeline can provide a potential solution to help researchers, decision makers, and medical doctors, among others, search and find the correct information in unique situation caused by the COVID-19 pandemic. We detailed the different components of this pipeline, including the traditional index-based information retrieval methods, the modern NLP-based neural network models, as well as insights and practical recipes to increase the quality of information retrieval of scientific publications targeted to the case of an infodemic. We grounded our results to the TREC-COVID challenge, where around 50 different teams participated in 5 rounds of competition. We showed very competitive results as judged by the official leaderboard of the challenge. Apart from the COVID-19 case, we believe our solutions can also be useful for other potential future infodemics.

## Acknowledgements

Innosuisse + CINECA

## Author contributions statement

D.T. conceived the experiments, D.T. and S.F. conducted the experiments, D.T., S.F., E.K. and P.A. analysed the results. N.B., E.K., D.V.A and P.A. prepared the data, S.F., D.T., J.C., R.G. and N.N. drafted the manuscript. All authors reviewed the manuscript.

## Additional information

### Competing interests

(mandatory statement).

The corresponding author is responsible for submitting a [competing interests statement](https://www.nature.com/srep/policies/index.html#competing) on behalf of all authors of the paper.

This statement must be included in the submitted article file.

## Footnotes

1 https://www.kaggle.com/allen-institute-for-ai/CORD-19-research-challenge

2 https://ir.nist.gov/covidSubmit

3 https://www.elastic.co

4 https://huggingface.co

5 https://github.com/jma127/pyltr

6 https://scikit-learn.org/stable/

7 https://github.com/usnistgov/trec_eval

